# Machine learning identifies signatures of host adaptation in the bacterial pathogen *Salmonella enterica*

**DOI:** 10.1101/204669

**Authors:** Nicole E. Wheeler, Paul P. Gardner, Lars Barquist

**Affiliations:** Wellcome Sanger Institute, Hinxton, United Kingdom; Biomolecular Interaction Centre, School of Biological Sciences, University of Canterbury, Christchurch, New Zealand; Department of Biochemistry, University of Otago, Dunedin, New Zealand; Institute for Molecular Infection Biology, University of Wuerzburg, Wuerzburg, Germany; Helmholtz Institute for RNA-based Infection Research, Wuerzburg, Germany

**Keywords:** hidden Markov model, random forest, loss-of-function, niche adaptation

## Abstract

Emerging pathogens are a major threat to public health, however understanding how pathogens adapt to new niches remains a challenge. New methods are urgently required to provide functional insights into pathogens from the massive genomic data sets now being generated from routine pathogen surveillance for epidemiological purposes. Here, we measure the burden of atypical mutations in protein coding genes across independently evolved *Salmonella enterica* lineages, and use these as input to train a random forest classifier to identify strains associated with extraintestinal disease. Members of the species fall along a continuum, from pathovars which cause gastrointestinal infection and low mortality, associated with a broad host-range, to those that cause invasive infection and high mortality, associated with a narrowed host range. Our random forest classifier learned to perfectly discriminate long-established gastrointestinal and invasive serovars of *Salmonella*. Additionally, it was able to discriminate recently emerged *Salmonella* Enteritidis and Typhimurium lineages associated with invasive disease in immunocompromised populations in sub-Saharan Africa, and within-host adaptation to invasive infection. We dissect the architecture of the model to identify the genes that were most informative of phenotype, revealing a common theme of degradation of metabolic pathways in extraintestinal lineages. This approach accurately identifies patterns of gene degradation and diversifying selection specific to invasive serovars that have been captured by more labour-intensive investigations, but can be readily scaled to larger analyses.

## Introduction

Understanding how bacteria adapt to new niches and hosts and thus emerge or re-emerge as a cause of infectious disease in human and animals is of critical importance to anticipating and preventing epidemic disease [1,2]. With the decreasing cost of genome sequencing, comparative genomics has become a rich source of insight into the origins and movement of bacteria in new pathogenic niches. However, translating whole genome sequence databases into mechanistic and functional insights remains a challenge.

Early expectations were that pathogen evolution would be driven primarily by the acquisition of virulence factors. However, as whole-genome sequencing has become increasingly routine, a decidedly more complex picture has emerged [3,4]. A pattern of bacterial entrance to a new niche followed by adaptation through the loss of antivirulence loci and reduced metabolic flexibility is now recognised as a paradigm of the emergence of important human pathogens from non-pathogenic bacterial species [5–8]. These new niches can be the result of virulence factor acquisition providing access to a previously inaccessible niche in a so-called foothold moment [8], or the emergence of new host niches driven by chronic disease [9–11]. While pathogen and host requirements for infection vary, there is increasing evidence of parallel evolution in bacteria adapting to the same or similar host niche. This is perhaps nowhere more evident than in the species *Salmonella enterica*.

*Salmonella enterica* strains that cause disease in warm-blooded mammals lie on a spectrum from those that have a broad host range and cause self-limiting gastrointestinal infection, to those that are more restricted in host range, but cause systemic disease and are typically associated with higher mortality [11,12]. Host-restricted, extraintestinal variants of *Salmonella enterica* have evolved independently multiple times from gastrointestinal ancestors [13], and show a greater degree of gene degradation compared to their generalist relatives [14–16]. There are common patterns in the genes that undergo pseudogenization in invasive *Salmonella*, most obviously an extensive network of genes required for anaerobic metabolism in the inflamed host [17,18], a pattern with parallels in other host-adapting enteropathogens [5].

Identifying these signals of parallel evolution has been challenging, relying mainly on manual annotation and comparison of pseudogenes [17,18]. Detection of pseudogenes in particular relies on ad-hoc criteria to identify large truncations, deletions, or frameshifts [19,20]. It is rare that the same genes or complete pathways are pseudogenized in host-adapted species; rather interpretation has relied on identifying overrepresentation of independent pseudogenization events clustered in certain pathways [17]. If pseudogenization leads to pathway attenuation or inactivation, it seems likely that reduced selective pressure will lead to a higher incidence of detrimental mutation fixation in other genes in these pathways. Indeed, we have previously shown that functional variant calling, based on sequence deviation from patterns of conservation observed in deep sequence alignments, shows a similar functional signal in host-restricted *Salmonella enterica* serovar Gallinarum to pseudogene analysis [21], identifying a larger cohort of genes where constraints on drift appear to have been lifted during host-adaptation.

In previous work we developed DeltaBS, a profile hidden Markov model (HMM) based approach to functional variant calling [21]. The basic assumption of this approach is that variation in conserved positions of a protein sequence is more likely to affect protein function than variation in less conserved regions. This approach can integrate information about nonsynonymous mutations, indels, and truncations. We have previously shown that DeltaBS can successfully identify functional changes in genes that would be missed by standard pseudogene analysis [22], and that a subset of genes in host-adapted strains appear to accumulate large DeltaBS values [21]. Additionally, others have observed similar changes in DeltaBS distributions during adaptation of *Salmonella* to a single immunocompromised host [10]. We generally assume that a large DeltaBS value is indicative of a decay in protein function, however a modest increase in DeltaBS associated with a phenotype may instead be indicative of diversifying selection.

Here, we have leveraged these previous observations to identify signatures of mutational burden consistent with adaptation to an invasive lifestyle. We have developed a random forest classifier using delta bitscore (DeltaBS) functional variant calling [21] that can perfectly separate intestinal *Salmonella* serovars from host-adapted, extraintestinal serovars. We use random forest models because they perform well on datasets with few informative variables [23,24], and the decision tree structure they employ has the potential to detect functional relationships (i.e. epistasis) between genes [25,26]. They have been applied successfully in the past to predict microbial phenotype using gene presence/absence data [27], and SNPs already known to be associated with phenotype [28,29]. We show that these models produce interpretable signatures of host-adaptation, and furthermore that these signatures can be detected in strains of *Salmonella* associated with invasive disease in immunocompromised populations in sub-Saharan Africa.

## Results

### Constructing a random forest classifier for extraintestinal Salmonellae

The approach taken in this investigation is summarised in Fig 1, and described below. We built our model using a collection of genomes from well-characterised reference strains of gastrointestinal and extraintestinal *Salmonella* serovars (S1 Table), drawing on the extensive curation of orthology relationships performed by Nuccio and Bäumler [17]. These strains were originally characterised as “gastrointestinal” or “extraintestinal” based on common patterns of gene degradation, host restriction and clinical characteristics observed among the extraintestinal strains [17], and we have employed this same categorisation our analysis. We scored the functional importance of sequence variation by comparing the protein coding genes of each serovar to profile HMMs from the eggNOG database [30], designed to capture patterns of sequence variation typically seen in the protein coding genes of Gammaproteobacteria (see Methods).

**Fig 1.**
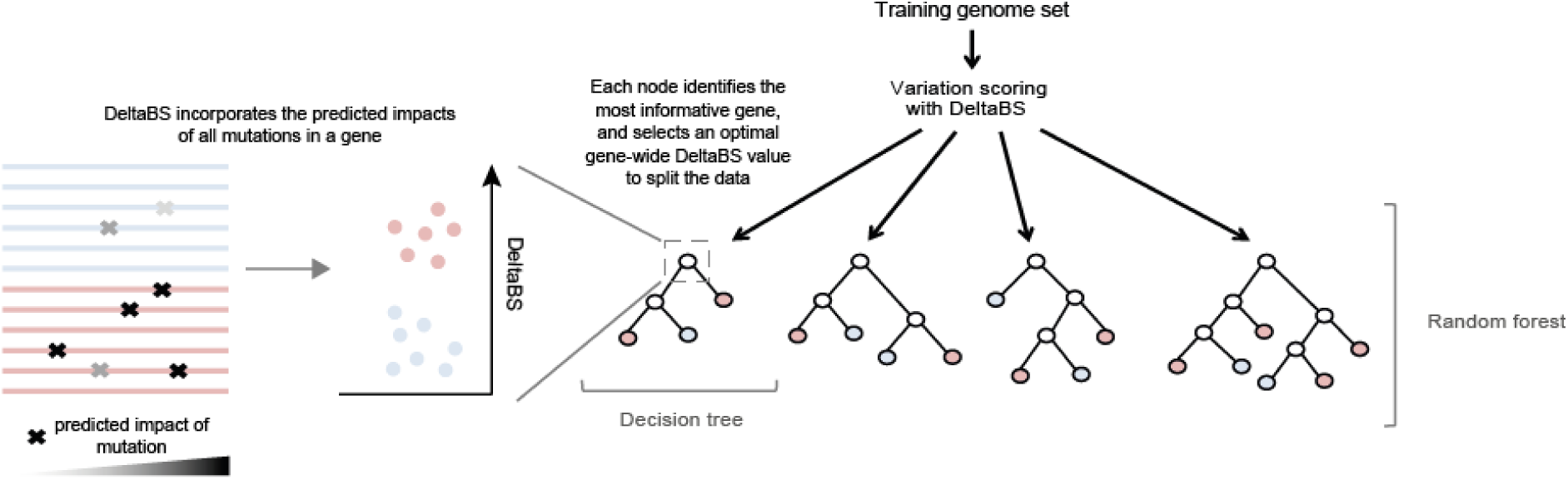
Overview of the approach employed in this study. For each genome, the functional significance of sequence variation within protein coding genes is quantified using the DeltaBS metric. Following scoring, a bootstrap sampling of genomes are used to train each decision tree. For each node in the tree, a random subset of genes are sampled, and the most informative gene from this set is chosen to split the data. For each node in the tree, the predictive utility of the selected gene (variable importance) is tested by calculating how well the gene separates the samples according to phenotype.

We then employed random forests to identify the genes which were most informative of phenotype when viewed collectively. Random forests work by building an ensemble of decision trees designed to predict a characteristic of the samples [31], in this case adaptation to an extraintestinal, or invasive, niche. For each node in the decision tree, the best gene of a random sampling from the training gene set is selected according to its ability to separate a randomly selected subset of samples by phenotype based on DeltaBS values. The process of building a random forest produces measures of variable importance that can be used to assess the relative utility of different genes in classification of *Salmonella* strains based on lifestyle.

### A small subset of genes are strongly predictive of invasiveness in Salmonella

To obtain an indication of the proportion of the genome that shows patterns of unusual sequence variation associated with an invasive phenotype, we trained a random forest model on a set of 6,438 orthologous genes. Accuracy of the model was assessed using out-of-bag accuracy. This out-of-bag (OOB) measure of accuracy gives us an indication of how well each decision tree in the forest performs at predicting phenotype in a serovar it has never encountered before, using information on DeltaBS differences collected from other serovars. Next, we performed iterative feature selection to improve the performance of the model. This process involved repeated rounds of selecting the top 50% of predictors and re-training the model, until the model achieved perfect OOB predictive performance on the training dataset (Fig 2A). When the full set of filtered orthologous genes was used to build a model, a subset of genes ranked much higher than the others in variable importance (VI) (Fig 2B). We then saw a tailing off of VI, resulting in 4,721 orthologous groups either not being used in the model, or not improving classification accuracy (as indicated by VI = 0). This set of genes was discarded in the the first round of feature selection, and a subsequent 1,521 genes were discarded in the subsequent three rounds. The final model used 196 of the original 6,438 genes for prediction (S2 Table). This model additionally achieved perfect classification accuracy on an independent set of genomes of the same serovars as our training data (Fig S1). We tested for overfitting using permutation tests, and for correlation bias [32] using a variety of alternative model building strategies, and found no evidence for either phenomenon in our model (File S1).

**Fig 2.**
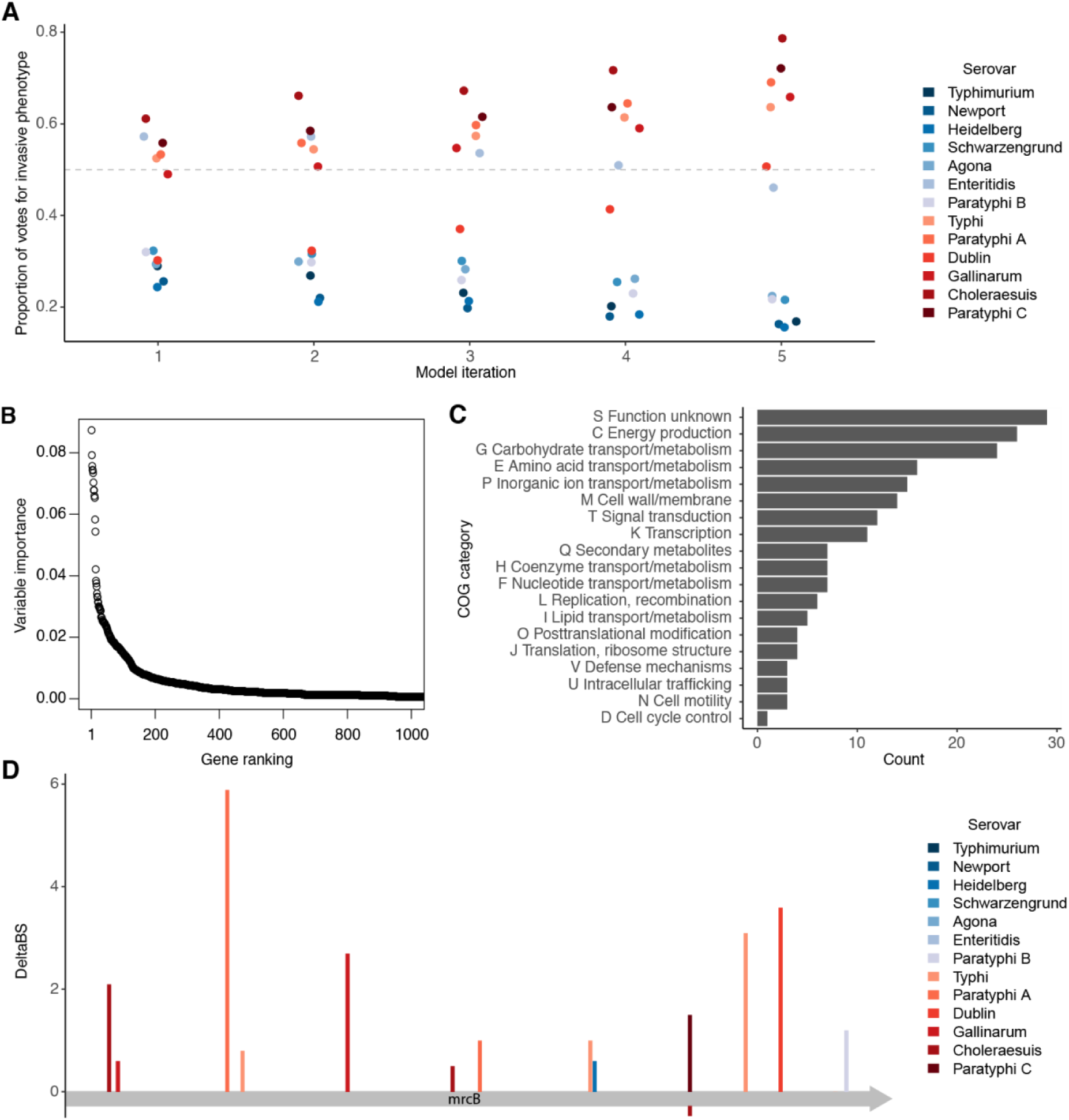
A subset of *Salmonella* genes are strongly indicative of invasive potential. A: Out-of-bag votes for phenotype of each serovar cast by each model. Model 1 is the model built using all predictor variables, then each successive model was built using sparsity pruning from the previous model’s predictor variables. Model 5 is the final model with 100% accuracy. Out-of-bag votes include only those votes cast by trees that were not trained on a given sample. The dashed grey line indicates the voting threshold to classify an isolate as invasive. Invasive serovars are coloured in red and gastrointestinal serovars are coloured in blue. B: Of all genes used in the original training dataset, a small minority are given high importance in identifying invasive strains. Variable importance is shown for the top 1000 genes used in the original training set. Variable importance was measured as average decrease in Gini index in a random forest model trained on all orthologous groups that met the inclusion criteria (N = 6,438). C: Functional categories associated with the top predictive genes. D: Mutations in *mrcB* (penicillin-binding protein 1b), one of the top three predictors. Mutations in different strains are colour-coded, with bars in red indicating a mutation in an extraintestinal strain and bars in blue indicating a mutation in a gastrointestinal strain. An estimate of the effect of the mutation on protein function (DeltaBS) is shown on the y-axis, with positive values indicating higher chance of a mutation impacting protein function. The x-axis represents the length of the protein.

### Predictive genes are typically degraded or absent in invasive isolates

We anticipated that the majority of informative genes identified in our study would be genes that showed functional degradation in invasive isolates but not in gastrointestinal isolates. Of the top predictors in our study (N = 196), 154 showed significantly greater mutational burden in extraintestinal strains compared to gastrointestinal strains (Mann-Whitney U test, adjusted *P*-value < 0.05), compared to 9 genes that showed significantly greater mutational burden in gastrointestinal strains. Of the genes that were more conserved in invasive isolates, one was the aldo-keto reductase *yakC*, which was deleted or truncated in all but one gastrointestinal strain and intact in all invasive strains. Another was the chaperone protein *yajL*, which appears to be important for oxidative stress tolerance [33,34].

Among the top predictors were several sets of genes belonging to the same operon (S2 Table). Examples included the *ttr, cbi* and *pdu* operons, which are all required for the anaerobic metabolism of 1,2-propanediol [35]. These operons have previously been identified as key degraded pathways in invasive isolates [16–18], and indicate the agreement of this method with other studies linking loss of gene function to host niche. Overall, a large proportion of the identified genes were involved in metabolism (Fig 2C), consistent with the findings of similar studies [17,18]. Of the 167 central metabolism genes identified by Nuccio and Bäumler (2014) as truncated or deleted in at least one extraintestinal serovar, only one of these was previously reported to be truncated in > 4 serovars. In contrast, we found that 20 of the 167 central metabolism genes were identified by our model as informative of phenotype, indicating that including signal from more subtle forms of loss of function improves our ability to detect parallelism across lineages of invasive *Salmonella*. Of the 13 genes reported to be frequently disrupted by Nuccio and Bäumler, our approach identified 9. The other 4 were either not a match to profile HMMs in our database, or the truncation did not fall within the span of the model. Other major categories affected include proteins involved in cell wall and membrane function, perhaps suggesting changes affecting recognition by the host immune system, and signal transduction, suggesting some degree of consistent regulatory rewiring during adaptation to an extraintestinal niche.

Information provided by multiple genes was often more informative of phenotype than a single gene individually, as was the case for *fimD* and *fimH* (Fig S2). FimD and FimH constitute central components of type 1 pili, and both are required for expression of normal fimbriae [36]. This demonstrates that our approach is capable of identifying epistatic relationships between genes, where a modification in function of one gene masks the functional status of the other.

### Sequence changes in key indicator genes involve independent mutations in each serovar, contributing to similar functional outcomes

When examining individual genes that showed differences in mutational burden between invasive and gastrointestinal isolates, we found that most of these mutations had occurred independently, and had occurred at different sites in the protein. Using a permissive threshold (DBS>3), or a conservative threshold (DBS>5), there were close to twice as many deleterious, independent mutations in the genes of the invasive serovars than those of the gastrointestinal (476:910; 537:991, respectively, see Methods). This phenomenon was even more pronounced when only mutations with DBS over the upper quartile were counted (249:612, Table S3). While the majority of genes identified appeared to be cases of gene degradation in invasive lineages, some genes showed more subtle signs of mutational burden, restricted to nonsynonymous changes of modest predicted functional impact.

An example of this, Fig 2D, illustrates mutation accumulation in one of the top candidate genes, *mrcB*, encoding penicillin-binding protein 1b (PBP1b). Not only does *mrcB* carry more mutations in invasive serovars compared to gastrointestinal serovars, the mutations have occurred independently in different positions within the protein. Penicillin-binding proteins are the major target of β-lactam antibiotics and are important for synthesis and maturation of peptidoglycan [37]. PBP1b in particular extends and crosslinks peptidoglycan chains during cell division. While PBP1b is not essential, it has been shown to be synthetically lethal when the partially redundant *mrcA*/PBP1a is deleted, and is important in *E. coli* for competitive survival of extended stationary phase, osmotic stress [38], and — in *Salmonella* Typhi — growth in the presence of bile [39]. Bile is an important environmental challenge for *Salmonella*, particularly for extraintestinal serovars which colonize the gall bladder [40]. While there are more mutations in invasive than in gastrointestinal serovars, the mutations that occur in this protein are all amino acid substitutions of modest predicted impact. This suggests that sequence changes could result in a modification of protein function, rather than a loss, consistent with the importance of PBP1b for the survival of *S*. Typhi during a typical infection cycle [39].

### S. Dublin and S. Enteritidis serovars are more difficult to classify than others

To anticipate the performance of our random forest model on new data we computed out-of-bag (OOB) error. Because random forests train each decision tree on a random subset of the training data, OOB error can be computed by testing the performance of these trees on data they have not been trained on, providing inbuilt cross-validation [31]. In our case, perfect OOB classifications were only achieved by the fifth iteration of the model. The need for iterative improvement of the model came from difficulty in correctly classifying the reference strains for serovars Enteritidis and Dublin. This is reflective of their relatively recent divergence and niche adaptation compared to other serovars in the study (Fig S3, Langridge et al. 2015). *S.* Gallinarum was classified much more readily than *S*. Enteritidis and *S.* Dublin, despite being closely related to both serovars, perhaps due to its host restriction.

*S.* Enteritidis was initially mis-classified as invasive, indicating that it shares genomic trends with invasive lineages. Genomic analyses have indicated that the ancestor of *S*. Enteritidis previously possessed intact pathogenicity islands (SPI-6 and SPI-19), each encoding a type six secretion system [18,41]. These loci have been implicated in host-adaptation and survival during extraintestinal infection [42,43], and it has been speculated based on their loss and other evidence that classical *S*. Enteritidis has been adapting towards greater host generalism with respect to its ancestral state [18]. This could explain the greater number of disrupted and deleted genes relative to other gastrointestinal serovars used in this study, and the difficulty in classifying it correctly. Conversely, *S*. Dublin was initially mis-classified as gastrointestinal. In previous studies *S.* Dublin has been shown to possess fewer pseudogenes than related invasive isolates [17,18], suggesting a lower degree of host adaptation than other invasive isolates. Indeed, *S*. Dublin is more promiscuous in its host range, primarily infecting cattle [44] while still causing sporadic human disease [45]. It seems likely that a subset of informative genes identified in early iterations of the model may have been indicators of host restriction or generalism rather than broad extraintestinal adaptation.

### Patterns of gene degradation identified in established invasive lineages are present in novel lineages of S. Typhimurium and S. Enteritidis associated with systemic infection

In recent years there have been reports of novel *S.* Typhimurium and *S.* Enteritidis lineages associated with invasive disease in sub-Saharan Africa [46–48] in populations with a high prevalence of immunosuppressive illness such as HIV, malaria, and malnutrition [49]. These lineages contribute to a staggering burden of invasive non-typhoidal salmonella (iNTS) disease, which is responsible for an estimated 3.4 million cases and circa 680,000 deaths annually [50]. Based on epidemiological analysis, high-throughput metabolic screening of selected strains, and analysis of pseudogenes it has been suggested that these lineages may be rapidly adapting to cause invasive disease in the human niche created by widespread immunosuppressive illness [11,46–48,51].

Two iNTS-associated lineages have recently been described within serovar Enteritidis [48], geographically restricted to West Africa and Central/East Africa, respectively. Initial observations have demonstrated that a representative isolate of the Central/East African clade has a reduced capacity to respire in the presence of metabolites requiring cobalamin for their metabolism and has lost the ability to colonize a chick infection model [48], suggesting adaptation to a new host niche. Similarly, two iNTS disease associated lineages have been described in serovar Typhimurium [47], both members of sequence type 313 (ST313), generally referred to as Lineage I and II in the literature. Lineage II appears to have largely replaced Lineage I since 2004, and it has been suggested this is due to Lineage II possessing a gene encoding chloramphenicol resistance [47]. Laboratory characterization of Lineage II strains has shown that they are not host-restricted [52,53], but do appear to possess characteristics suggestive of adaptation to an invasive lifestyle [54–57], though it is important to note that this is a complex trait and not easily quantified.

Given the evidence of adaptation to an invasive niche in these lineages, we asked if genomics signatures of extraintestinal adaptation we had detected previously could be detected in iNTS disease associated lineages. To this end, we applied our predictive model trained on well-characterized extraintestinal strains to calculate an invasiveness index, the fraction of decision trees in the random forest voting for an invasive phenotype. First, we compared isolates from African iNTS-associated clades of *S*. Enteritidis (N=233) to a global collection of isolates generally associated with intestinal infection (N=100) [48].

Our model gave iNTS-associated *S*. Enteritidis strains a higher invasiveness index than the globally distributed isolates (Fig 3A,B, Table S4), indicating the presence of genetic changes paralleling those that have occurred in extraintestinal serovars of *Salmonella*. Similar gene signatures were only rarely observed in the global epidemic clade (Fig 3C). These findings are consistent with the metabolic changes observed by Feasey et al. [48] in the Central/Eastern African clade compared to the global epidemic clade. In particular we found signs of gene sequence variation uncharacteristic of gastrointestinal *Salmonella* across a number of key genomic indicators, including *tcuR, ttrA, pocR, pduW, eutH*, SEN2509 (a putative anaerobic dimethylsulfoxide reductase) and SEN3188 (a putative tartrate dehydratase subunit), all in pathways previously identified by Nuccio and Bäumler [17] as being involved in the utilization of host-derived nutrients in the inflamed gut environment. This indicates that our model is able to identify early signatures of adaptation, even in these recently emerged strains that still retain some capacity to cause enterocolitis [48].

**Fig 3.**
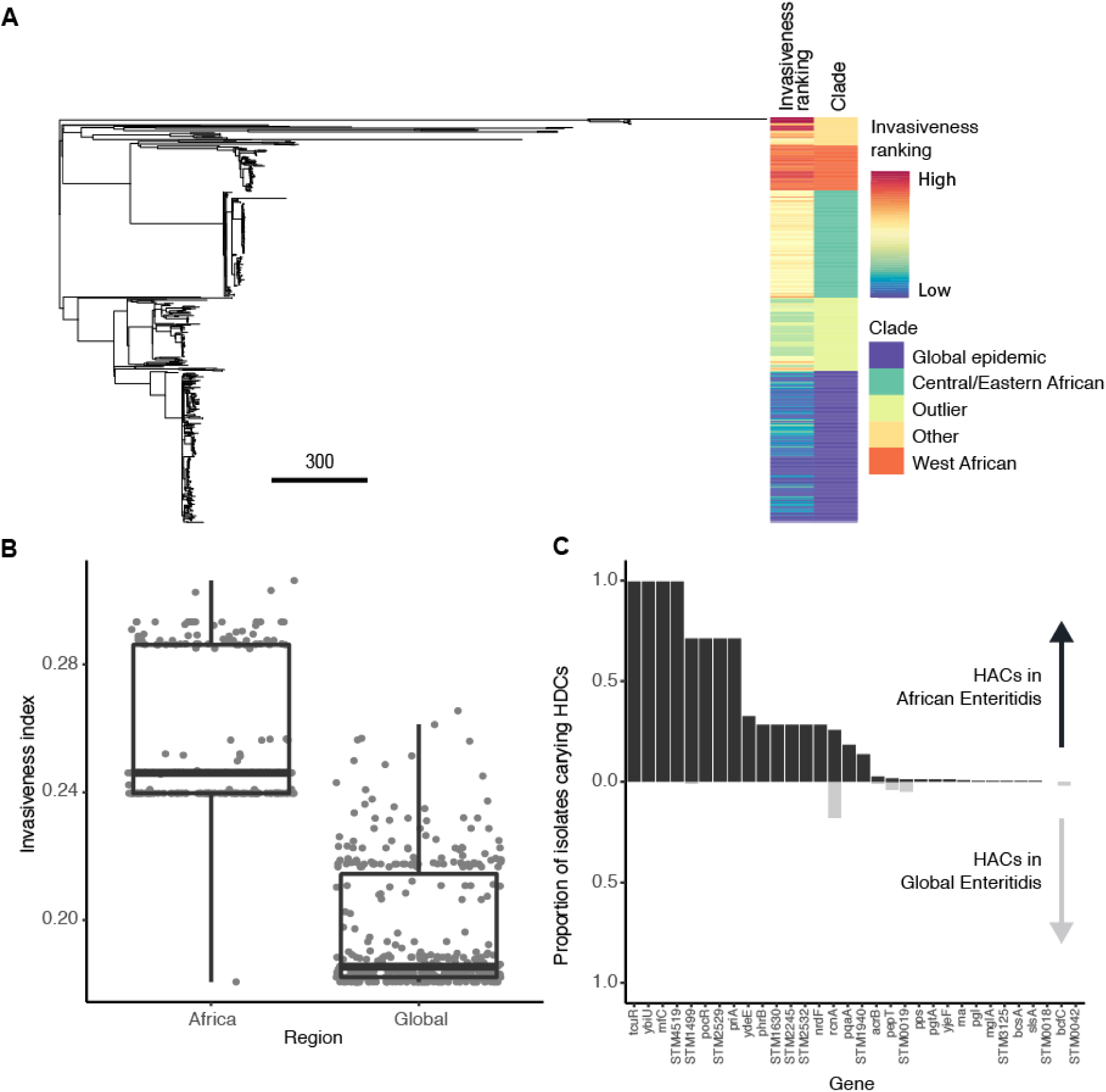
Voting of the model on African iNTS and global gastrointestinal isolates. A: Maximum likelihood phylogeny of all *S*. Enteritidis isolates included in the study, annotated with invasiveness ranking and clade (note: Outlier refers to the distinct sister clade of the global epidemic strains identified by [48], while Other refers to strains that don’t belong to a named clade). B: Invasiveness indices for African and non-African clades of *Salmonella*. Lower and upper boundaries of the boxplots correspond to the 25th and 75th quantiles. C: The proportion of isolates from each tested dataset carrying a hypothetically attenuated coding sequence (HAC, defined by a DeltaBS>3 relative to the reference serovar). Genes are ordered by the amount of degradation observed in African clades. African strains are shown in the positive y-axis in darker grey, global strains are shown in the negative y-axis in lighter grey.

To confirm this, we performed an additional comparison of *S*. Typhimurium ST313 isolates (N=208), to global isolates from other STs, predominantly ST19, associated with gastroenteritis (N=51) [51,58]. Similarly to iNTS associated *S*. Enteritidis isolates, *S*. Typhimurium ST313 isolates has a higher invasiveness index than isolates from other STs (Fig S4, Table S5). Within ST313, Lineage II scored higher than Lineage I, possibly suggesting differential adaptation to the extraintestinal niche. We found that there were in fact more degraded genes unique to Lineage I than Lineage II, but that these genes were assigned less weight in the model, so did not impact score as strongly (Figure S2 & S3). Interestingly, ST313 has recently been shown not to be entirely restricted to Africa, with isolation reported in Brazil [59] and the UK [58], associated primarily with gastrointestinal disease. We included a collection of UK ST313 strains [58] in our analysis, and found that their invasiveness index tended to be elevated compared to non-ST313 salmonellae, and intermediate between Lineage I and II, suggesting that this adaptation is not restricted to circulating African strains, as it can be seen in strains collected from other countries as well (Fig S5). This observation is consistent with the work of Ashton et al., who noted shared pseudogenes and phenotypic traits in UK and African ST313 isolates. This suggests our model is capturing features here associated with the ability to colonize an extraintestinal niche, rather than enter it in healthy individuals.

In addition to the iNTS lineages we investigated, some other strains had unusually high invasiveness indices. Among the top scoring isolates outside of the African *S*. Enteritidis lineages are Ratin strains, a rodenticidal lineage used as commercial rat poison before the 1960s [60]. In *S*. Typhimurium, a clade containing strains DT99, DT56 and U313 also scored highly. These strains appear to be adapted to birds, and DT99 and DT56 have been reported to be highly virulent in pigeons [12,61–63].

While the above data suggests that our model is detecting genetic changes associated with extraintestinal survival, it is difficult to infer directionality from large isolate collections. We have addressed this using a unique case of accelerated adaptation over the course of a single infection (Fig 4). We scored the invasiveness index of a collection of hypermutator *S*. Enteritidis isolates collected over a ten year period that were adapting to chronic systemic infection of an immunocompromised patient [10]. We found a significant positive correlation between invasiveness index and duration of carriage (r=0.96, n=6, *P*=0.002). Additionally, there was a significant shift over time in the DeltaBS distribution for the genes in our model as compared to the rest of the genome (*P*=7.576e-05, Mann Whitney U test). This suggests a specific change in selective pressure on genes inferred to be important for extraintestinal survival from established invasive serovars, and provides evidence for parallel adaptation.

**Figure 4.**
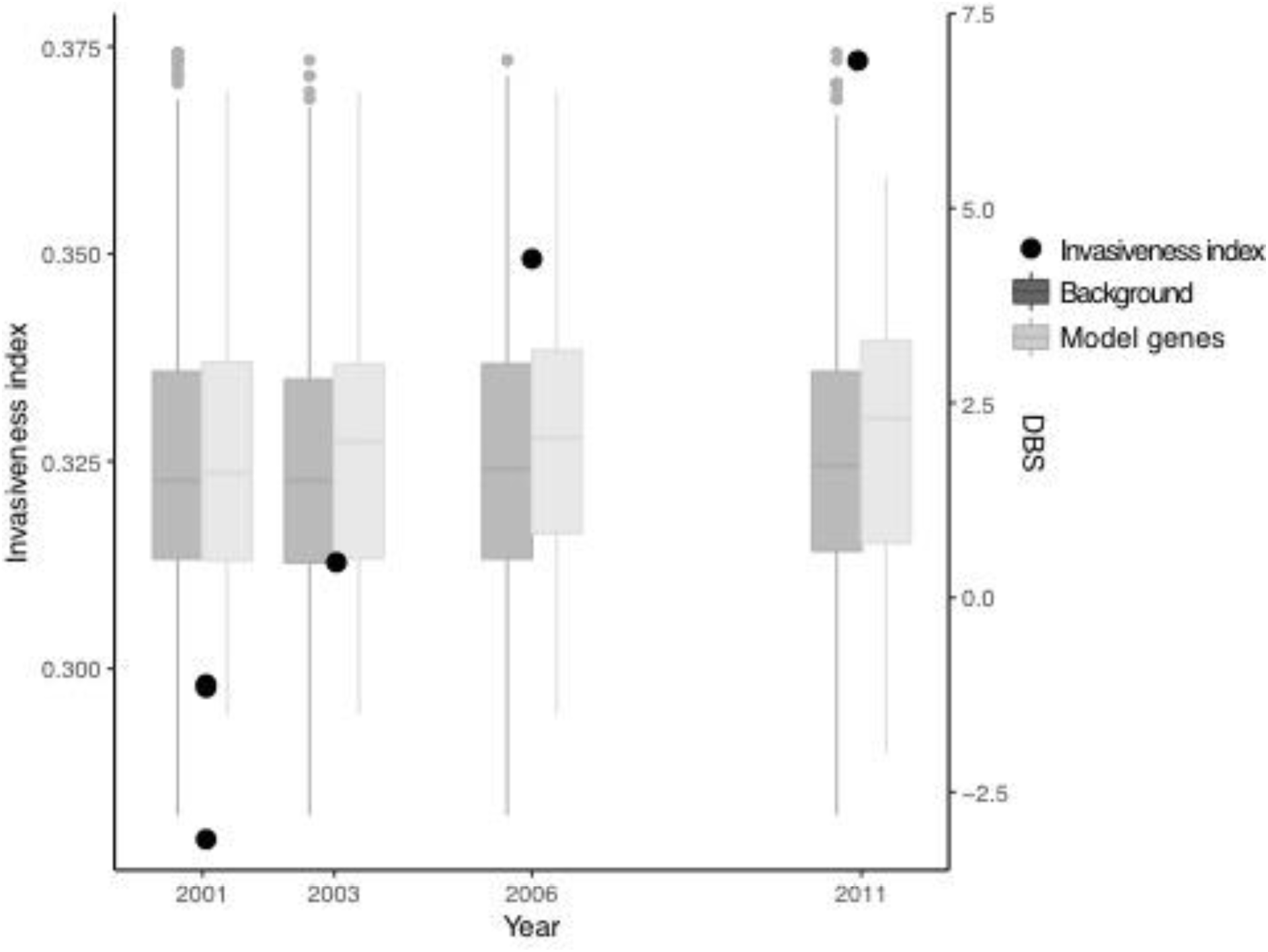
Invasiveness indices and DeltaBS (DBS) values for isolates collected during long term invasive infection of an immunocompromised patient (Klemm et al. 2016). Black points show the increase in the invasiveness index over time. Boxplots show a significant shift in DBS distribution over the duration of carriage for genes selected by our model built from well-characterised invasive serovars as compared to the rest of the proteome. Isolates from [10]. DBS distributions for 2001 have been pooled, but are representative for all three isolates individually. The y-axis for DBS values has been truncated for better visualisation.

## Discussion

Parallel evolution appears to be common in niche adaptation, which allows us to identify genes that are important for survival in different environments [64]. Parallelism has been observed across vastly different time scales in adapting pathogens. Parallel evolution in the distantly related genuses *Salmonella* and *Yersinia* during adaptation to invasive infection of the human host has led to independent losses of the *ttr, cbi* and *pdu* genes, important for anaerobic metabolism during intestinal infection [5]. Within genuses, parallelism has been observed when distinct lineages acquire similar virulence factors leading to similar phenotypes, as with *Yersinia pseudotuberculosis* and *enterocolitica* [8], or the repeated emergence of the *Shigella* phenotype within the *Escherichia* [6]. Even on the scale of a single human lifetime, parallel adaptation has been observed in *Pseudomonas aeruginosa* lineages adapting to infection of the lungs of children with cystic fibrosis [9], or a hypermutator strain of *Salmonella* adapting to an immunocompromised host [10]. With pathogen sequencing for disease surveillance becoming increasingly routine [65–67], we have the opportunity to search for signals of parallel evolution as new pathogens emerge, or old pathogens expand into new niches.

Here, we have developed an approach for automatically learning which genes contribute to this parallel adaptation. Leveraging the DeltaBS functional variant scoring approach we developed previously [21] allowed us to construct scores which integrate independent mutations and indels that impact gene function. Using these scores, we were able to construct a classifier model which is able to separate *Salmonella* serovars adapted to an extraintestinal niche from gastrointestinal strains. Importantly, the random forest classifier that we used produces interpretable lists of genes involved in this adaptation, which agree with results in the literature attained through manual curation of pseudogenes. Additionally, we have shown that this classifier is able to identify nascent signatures of adaptation in strains of *Salmonella* which have been evolving in response to large populations of immunocompromised patients in resource-poor nations.

Other automated approaches to detecting adaptation have been developed which search for SNPs [68] or words [69,70] associated with phenotype. These approaches, termed microbial genome-wide association studies (GWASs), have used techniques adapted from human GWASs, but better cater to methodological issues that arise due to the differences between human and bacterial inheritance patterns. Major differences impacting analyses are stronger linkage disequilibrium (LD) between genetic variants in bacterial genomes, greater population stratification, and often stronger selection for traits [71]. Greater LD and population stratification often result in traits being linked closely with particular lineages, and a large number of variants unique to a lineage being spuriously associated with phenotype. Correction for population stratification allows greater discrimination of true and false positive associations, but results in a substantial loss of power to detect true positives [71], particularly in phenotypes that are highly polygenic and are not under strong positive selection [72]. This can be corrected by increasing the sample size of the study, but increasing sample size can make measurement of complex phenotypes infeasible [23].

A number of machine learning approaches to predicting phenotype from genotypic information have also been recently developed. A notable example is a Support Vector Machine (SVM) based approach to predicting host range in *Salmonella enterica* and *Escherichia coli* [73], as it has a similar aim of predicting strains with a higher probability of causing severe disease. We have taken a markedly different approach to other machine learning based studies, primarily in our use of few, distantly related training examples, rather than densely sampled strains across a narrower phylogenetic distance. This is because we wanted to prevent over-fitting of the model through the inclusion of predictors that were informative of phylogeny rather than phenotype, and we wanted an accurate estimation of predictive error, which cannot be achieved using traditional cross-validation when there is a strong correlation structure in a dataset [74]. We have also taken additional steps to examine the genes and criteria used by the model to make predictions, and have presented these in Supplemental Table S2, in order to aid the reader’s understanding of how the model makes predictions, and what this teaches us about the biology of this phenotype.

The use of DeltaBS output as training variables differs from current approaches by allowing the estimation of the combined effects of variants, both common and rare, on gene function. The weighting scheme can also combine data on gene presence/absence, indels and SNPs into a single metric. It significantly reduces the number of association tests that need to be performed to comprehensively capture much of the genetic diversity in a species, increasing power to detect associations, and reducing the requirement for such large sample sizes. The approach also aids in identifying genetic variants that are most likely to have a phenotypic effect within LD blocks. The DeltaBS variant scoring approach can be readily applied to large datasets, and could be employed in a linear mixed model (LMM) based association testing framework [68], or used in a hybrid LMM-random forest based approach [75] to preserve the ability of the metric to detect epistasis between genes [26].

## Conclusions

In this study, we have demonstrated the insight to be gained by the layering of machine learning approaches to better understand niche adaptation in a bacterial pathogen. Firstly, profile hidden Markov models allow us to capture information on common patterns of sequence variation in protein families in order to understand the functional significance of specific mutations. Using data on the accumulation of functionally impactful mutations across the proteome as input, random forests then allow us to identify genes that display a difference in selective pressures between lineages with different phenotypes. Not only has this approach proved effective at identifying biological mechanisms behind bacterial niche adaptation, it has also allowed us to detect the emergence of new extraintestinal lineages by searching for these recurrent patterns of mutation accumulation in a way that allows the recognition of novel mutations as cases of the same underlying shift away from the sequence constraints a gene is usually subjected to. We believe this general approach will be broadly applicable to any pathogen where multiple lineages are adapting to the same niche, and will be able to detect signatures of adaptation that are missed by other methods.

## Methods

### Genome data and identification of orthologs

High quality genomes for 13 well-characterised *Salmonella enterica* serovars were retrieved from the NCBI database (accessions and serovar information can be found in S1 Table). The serovars were divided into gastrointestinal and extraintestinal serovars according to the classifications made by Nuccio and Bäumler [17]. Ortholog calls were also taken from the Supplementary Material of Nuccio and Bäumler [17]. A core gene phylogeny for the strains used to build the model was produced using RAxML [76], based on a core gene alignment created in Roary [77].

### Measuring the divergence of genes from predicted sequence constraints

Profile hidden Markov models (HMMs) for Gammaproteobacterial proteins were retrieved from the eggNOG database [30]. We chose this source of HMMs because it is publicly available, allowing for better reproduction of analyses, and we feel it provides a good balance between collecting enough sequence diversity to capture typical patterns of sequence variation in a protein, without sacrificing sensitivity in the detection of deleterious mutations, as we have observed with Pfam HMMs [21]. Each protein sequence was searched against the HMM database using hmmsearch from the HMMER3.0 package (http://hmmer.org). The top scoring model corresponding to each protein was used for analysis (N = 8,060 groups). Orthologous groups (OGs) with no corresponding eggNOG HMM, or more than one top model hit were excluded from further analysis (N = 1,524). If most genes in an OG had a significant hit (E-value<0.0001) to the same eggNOG model, any genes within this OG that did not were assigned a score of zero, reflecting a loss of the function of that protein. These cases typically reflected a truncation that had occurred early in the protein sequence. Additionally, genes with no variation in bitscore for the match between protein sequences and their respective eggNOG HMM across isolates were excluded (N = 188). After this filtering process, 6,439 orthologous groups remained for analysis. Residue-specific DeltaBS (as in Fig 2D) was calculated by aligning orthologous sequences, choosing a reference sequence (from *S*. Typhimurium), and substituting each variant match state and any accompanying insertions into the reference sequence and calculating the difference in bitscore caused by the substitution.

### Training a random forest classifier

The R package “randomForest” [78] was used to build random forest classifiers using a variety of parameters to assess which were best for accuracy. We used out-of-bag (OOB) error rate to measure the performance of the model [31]. Out-of-bag error is calculated automatically by the randomForest R package as the model is built. Briefly, calculations are performed as follows: as each decision tree is trained using a bootstrap sampling of the training genomes, a small number of samples are left aside to test the predictive accuracy of each decision tree on previously unseen samples. For each serovar, votes are collated and accuracy is calculated from only those decision trees that did not include the serovar in their training set. In this application, this step tests whether the genomic signatures of invasiveness captured by the decision trees based on some serovars are present in other serovars, and thus whether the model can detect adaptation to an invasive lifestyle in previously unseen lineages. OOB error rate, stabilised at 10,000 trees, so we chose this as a parameter for optimising the number of genes sampled per node (mtry). mtry values of 1, *p*/10, *p*/5, *p*/3, *p*/2 and *p* (where *p* = the number of predictors) were tested, and we found that at mtry=*p*/10, the number of genes that were either not incorporated into trees, or did not improve the homogeneity of daughter nodes when they were incorporated into trees (as measured by mean decrease in Gini index, [79]) stabilised at ∼92%. Training the random forest classifier over five iterations took 55 seconds on a laptop computer. In order to assess how well this method would scale, we trained another model on a larger dataset of *S*. Enteritidis strains (N=677) using the same workflow and site of isolation as a proxy for phenotype, which took 28 minutes.

To improve the performance of the model, we performed five model building and sparsity pruning cycles. For the first cycle, we built a random forest model using all genes that met the inclusion criteria, and performed sparsity pruning by eliminating all variables that had a mean Gini index (variable importance) of zero or lower (meaning the gene was either not included in the model or did not improve model accuracy when it was). Four successive rounds of model building and sparsity pruning involved building a new model with the pruned dataset, then pruning the genes with the lowest 50% of variable importances. The resulting model had 100% out-of-bag classification accuracy. We also tested the accuracy of the full model on a collection of alternative strains related to the training dataset (see Table S1). Orthologs to the top genes identified by our model were identified using phmmer from the HMMER3.0 package (http://hmmer.org). Additional notes on model building and testing are provided in File S1.

We tested the top 196 genes for the presence of independent mutations in each serovar by aligning each sequence to the profile HMM representing that protein family. Variation in each sequence with respect to a designated reference sequence from the set (as selected by Nuccio and Bäumler, 2014) at each site in the HMM was identified and classified as either a mutation unique to a single serovar, or one shared among mutliple serovars. Consecutive deletions or insertions with respect to the HMM consensus sequence were collapsed into single mutational events.

### Invasive non-typhoidal Salmonella analysis

Read data from Feasey et al. [48] and Klemm et al [10] was mapped to the reference genome *S*. Enteritidis P125109. Reads from Okoro et al. [51] and Ashton et al. [58] were mapped to the reference genome *S*. Typhimurium LT2. For samples in the Okoro study, if an isolate was sequenced using multiple runs, the most recent run was chosen for analysis. All reads were mapped using BWA mem [80] and regions near indels were realigned using GATK [81]. Picard (http://broadinstitute.github.io/picard) was used to identify and flag optical duplicates generated during library preparation. SNPs and indels were called using samtools v1.2 mpileup [82], and were filtered to exclude those variants with coverage <10 or quality <30. For tree building, a pseudogenome was constructed by substituting high confidence (coverage >4, quality >50) variant sites in the reference genome, and masking any sites with low confidence with an “N”. Insertions relative to the reference genome were ignored, and deletions were filled with an “N”. Pseudogenome alignments were then used as input to produce trees using Gubbins [83] to exclude recombination events, and RAxML v8.2.8 [76] to build maximum likelihood trees using a GTR + Gamma model. Samples with >10% missing base calls were excluded from the analysis.

Sequences for the 196 genes of interest used in the random forest model were retrieved for each isolate and translated. These were then scored using their respective profile HMMs. Score data was collated, and any missing values were marked as ‘NA’ and imputed using the na.roughfix function from the randomForest R package [78]. This is a different approach used to that of the training dataset, due to the potentially lower quality of the sequenced genomes leading to gene absence due to low coverage rather than true deletion or severe truncation. The relationship between invasiveness ranking and phylogeny were visualised using Phandango [84].

### Data availability

All genome sequence data are publicly available, and accessions are provided in the appropriate Supplemental Tables. Code and data for reproducing this analysis, performing an equivalent analysis using new data, and assessing the invasiveness index of other *Salmonella* strains is publicly available at http://www.github.com/UCanCompBio/invasive_salmonella.

## Acknowledgements

We are grateful to Sean Eddy for useful discussions and providing fast, accurate and free software, and to Simon Harris for developing the pipeline used for mapping reads and calling SNPs for the iNTS portion of our analysis. We also thank Julian Parkhill, Nick Feasey, Nick Thomson, Alexander Westermann, Stan Gorski, and John Crump for their helpful feedback.

