## Supplementary Materials for "Machine learning identifies signatures of host adaptation in the bacterial pathogen *Salmonella enterica*"

**Building and testing the random forest model**

**Choosing model building parameters**

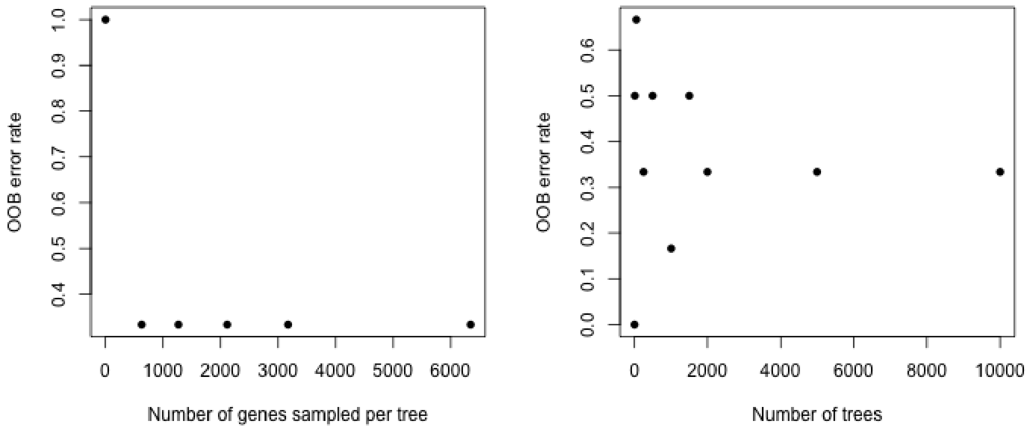


To select model training parameters, we used OOB error rate to evaluate a range of values. Error rate appeared to stabilise at 10,000 trees, so this value was used for ntree. We tried values of 1, n/10, n/5, n/3, n/2 and n for mtry. Error stabilised relatively early. We decided to use mtry=n/10 as it would reduce the chances of sampling correlated predictors.

**Permutation testing for over-fitting**

With many parameters and few samples, overfitting to random fluctuations in the data may be a concern. To address the risk of this given our dataset, we produced 1,000 phenotype permuted datasets and re-ran our model building pipeline (5 rounds of feature selection, ntree=10,000, mtry=n/10). We measured the accuracy of each model built using this approach, and compared the performance of our real model with the models built on randomised phenotypes. We found that the accuracy of our model after five iterations of feature selection (OOB accuracy = 1) was exceptional compared to models trained on permuted phenotypes, with only 5 permutations (P=0.005) attaining perfect OOB accuracy.

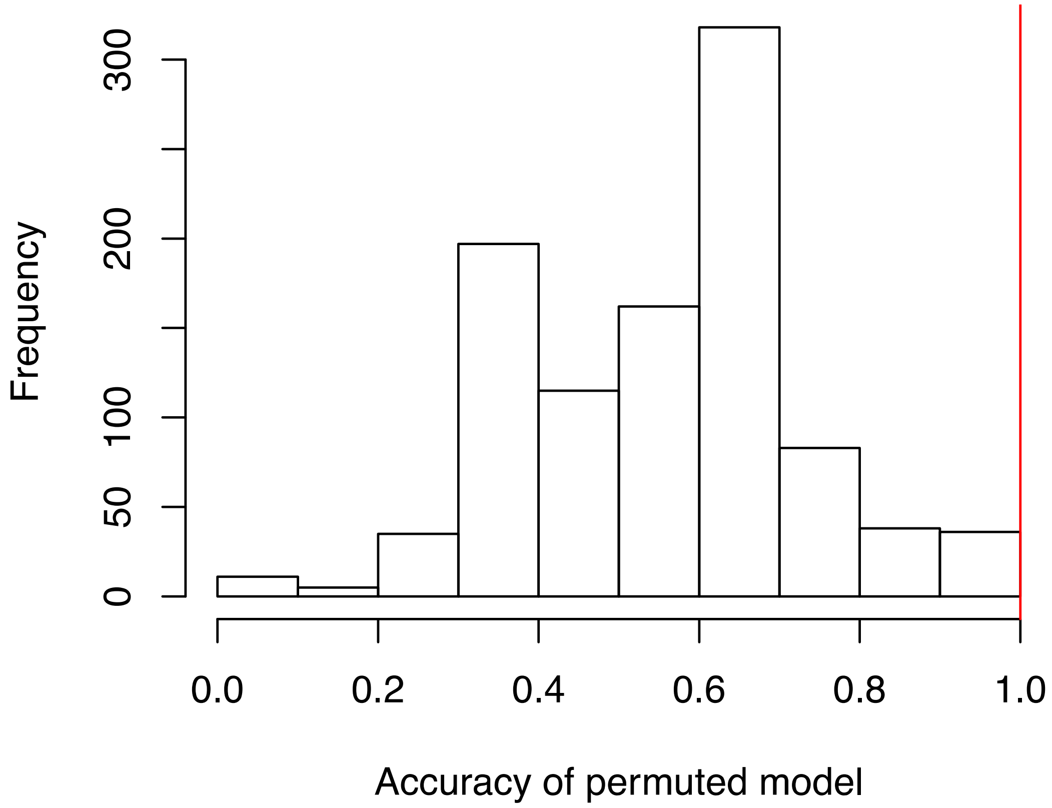

**Comparison of signal in real vs. randomized data**

To test for signal in the bitscore pattern found in real data (in other words, are there genuinely more bitscore patterns that are consistently different between our pathovars than by chance?), we permuted the values of each gene separately, then ran a single model-building step and looked at the distribution of variable importances, as measured by mean decrease in accuracy.

Across ten different permutations of the variable values, the real variable importances (VI), as measured by mean decrease in accuracy, were consistently higher overall than those for the permuted variables (Mann-Whitney U test $P<2.2E-16$). In addition, twice as many permuted values as real values were given negative VI values (~1000 vs ~500), a strong indicator that the variable has no relationship with phenotype. A representative distribution of real (red) and permuted (white) feature importances are shown below.

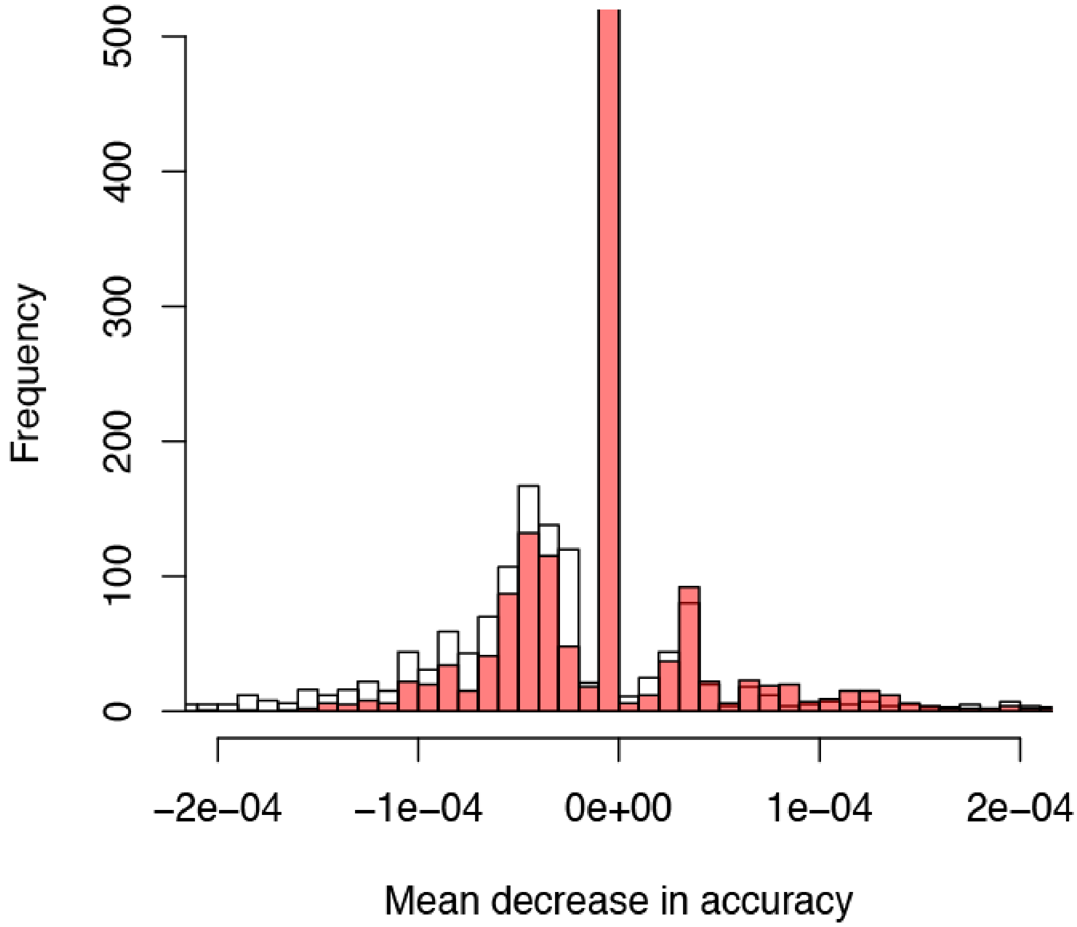

**Testing for correlation bias**

In order to test whether our final gene set was affected by correlation bias (i.e. whether informative genes were excluded because they were correlated), we applied a number of clustering approaches following the example of Tolosi and Lengauer "Classification with correlated features: unreliability of feature ranking and solutions", Bioinformatics 2011, then re-ran the same model building pipeline to check the overlap in our gene sets.

**Overview of approach**

As a general approach, after choosing a method of normalising the data, we computed a distance matrix and performed hierarchical clustering. We re-built the model using hierarchical clustering of bitscore values (using both a scaled and centered normalisation procedure and a rank-based procedure to account for outliers in our bitscore distributions). We clustered using either Euclidean distance (for scaled variables) or 1-Spearman correlation coefficient (for ranked), and took the centroid of each cluster as input variables for our model. We used either Gini index or mean decrease in accuracy as measures of variable importance to select variables at each iteration of the model-building process.

Following hierarchical clustering, we broke the gene sets into 500, 1000, 2000, 2500, 3000, 3500, 4000, 5000 and 6348 (one for each gene) clusters. We then ran model building using the same parameters as the original model (ntree=10000, mtry=n/10) for 6 iterations (the original model only took 5 iterations to reach 100% accuracy, but some of these approaches took longer). The first round of feature selection removes any clusters with VI=0, then subsequent rounds discard the lowest ranked 50% of clusters. Each condition was run twice, to account for variability in outcome from random sampling.

**Model types**

We have run each normalisation, distance method, and method of measuring VI through the iterative model building process for 6 iterations (one more than the original model). The following plots show:

1. out-of-bag accuracy
2. overlap with the original gene set
3. overall numbers of genes in the original model, the new model and both

to compare the approaches.
Plots are named according to the pattern:
normalisationMethod_distanceMeasure_variableImportanceMeasure

#### Accuracy of the models

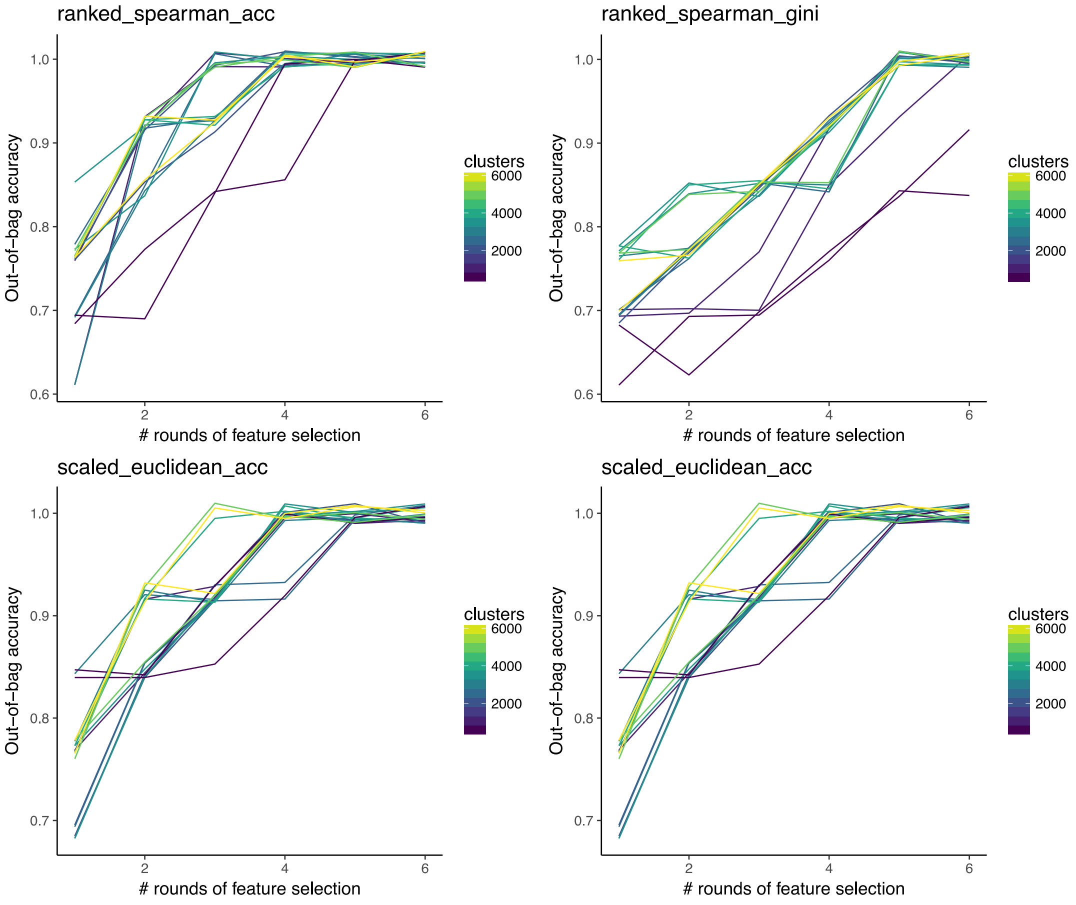

### Inclusion of the original 196 genes in the different types of model

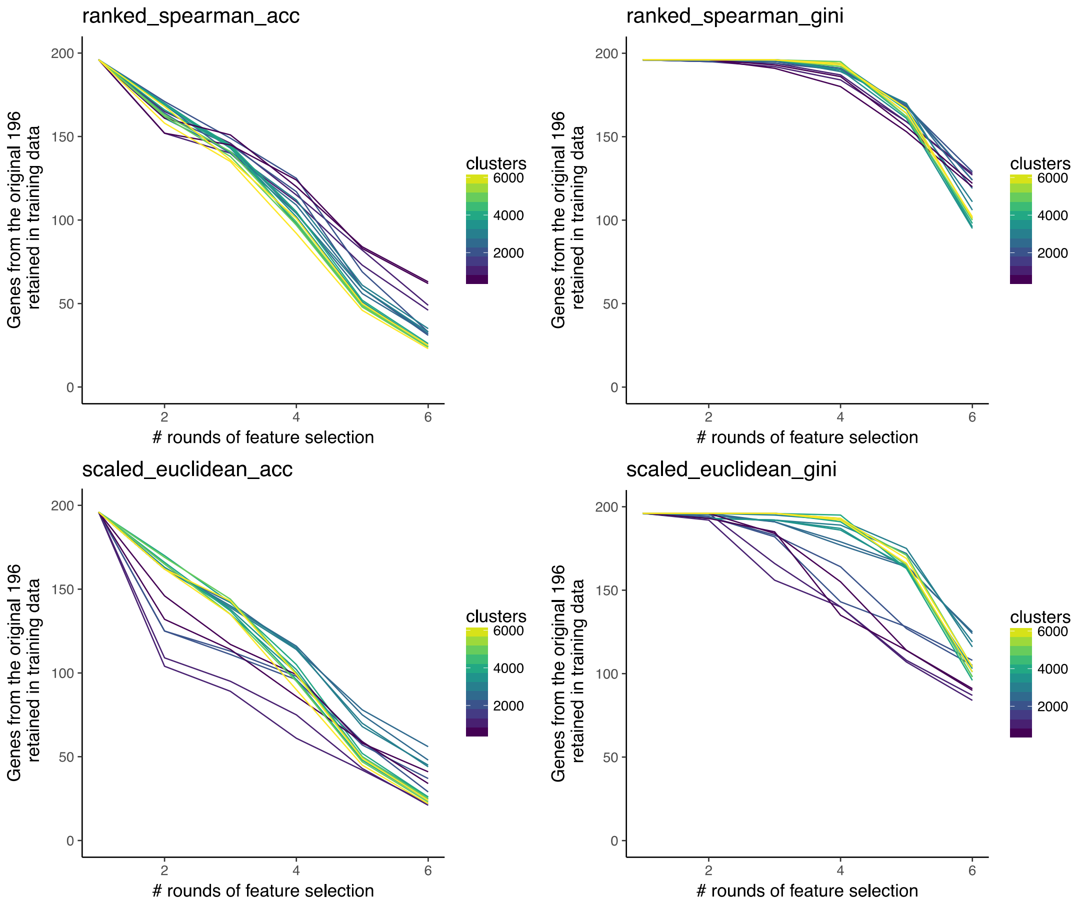

**Looking at overall gene sets included in the model**

Note: iteration 1 is not shown, since all genes are included in all models. This truncation of the y-axis was done in order to better view the smaller values. In general, all methods roughly recapitulate our initial gene set as they approach perfect accuracy, and become a subset of our gene set on further iteration.

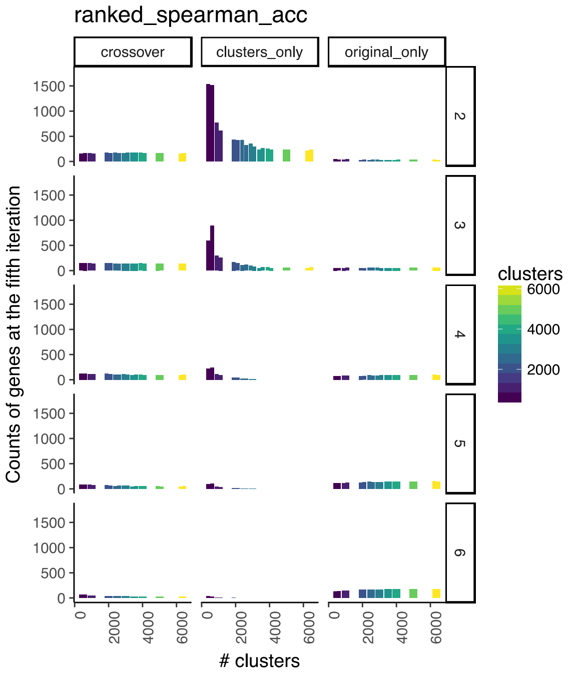


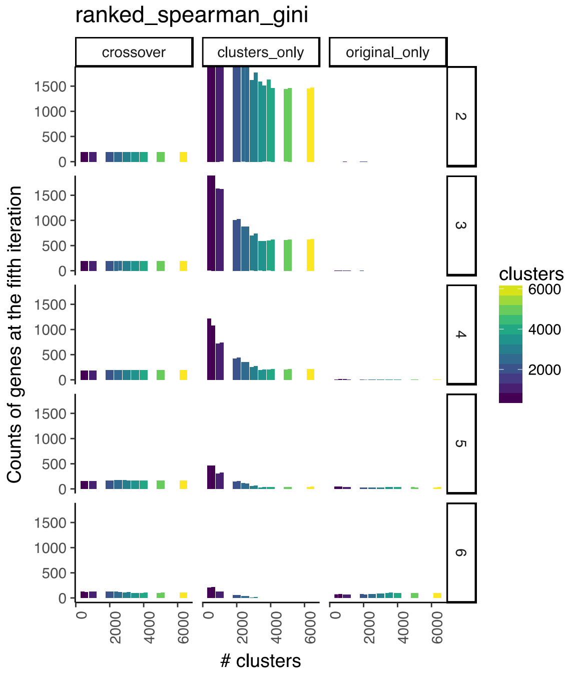


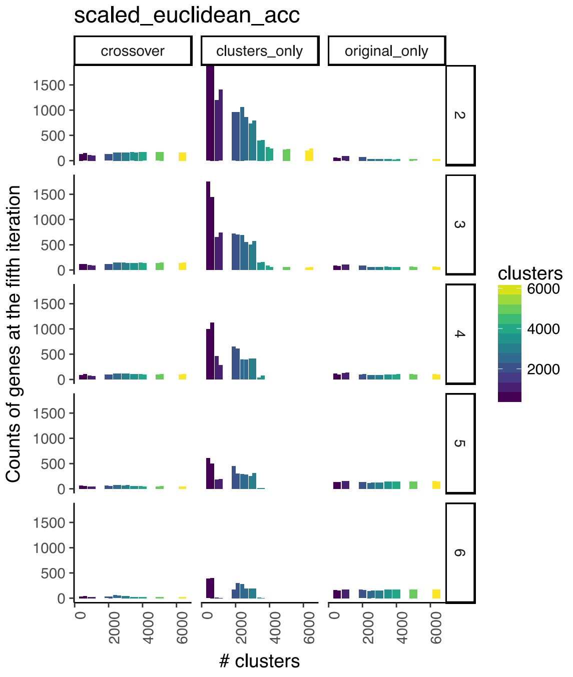


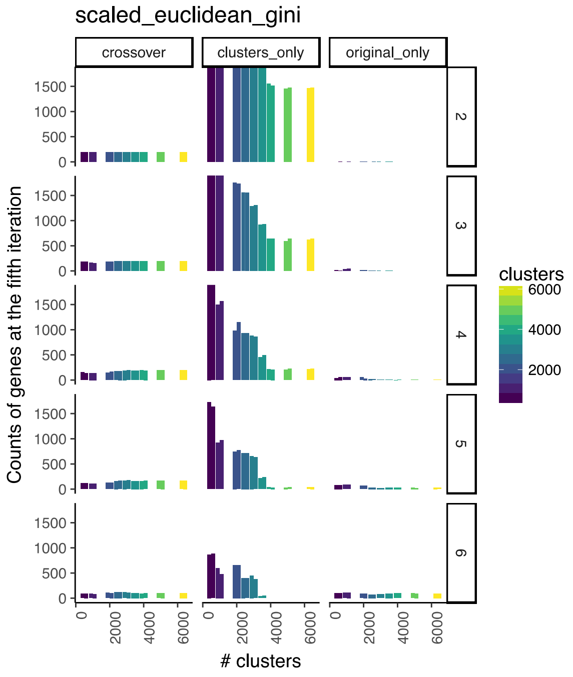

**Comments**

Out-of-bag accuracy seems to peak around 3-5000 clusters, depending on the metric used to cluster and choose genes. In this range, with models that have reached 100% accuracy, there are few or no additional genes selected by these methods that weren't included in our original gene set, indicating that our feature selection has not resulted in exclusion of informative genes due to correlation bias. We also observed that over later iterations, genes not included in our core set are discarded preferentially to our core gene set. As the number of clusters approaches the number of genes, the correlation between gene and cluster variable importance improves, to the point where there are no notable outliers over 3000 clusters for rank-based clustering and over 3500 for clustering of scaled, centred values.

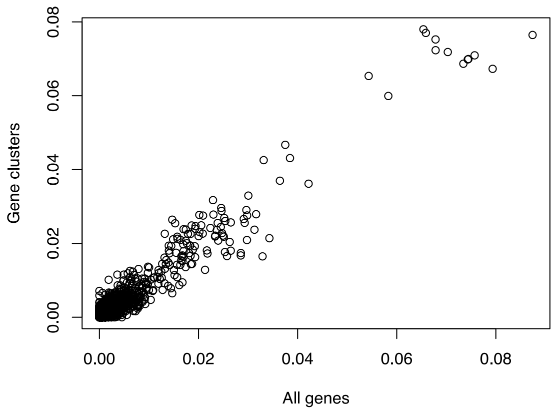


Ranked Spearman Gini, 3500 clusters

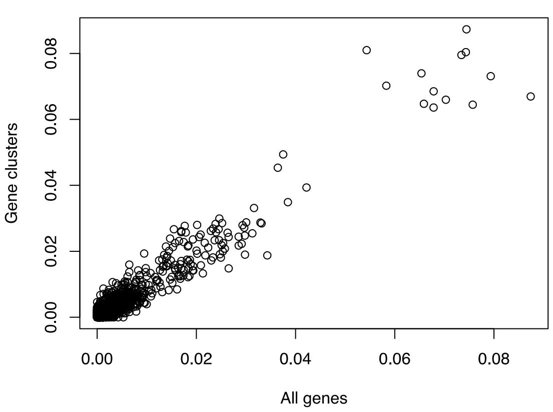


Scaled Euclidean Gini, 4000 clusters

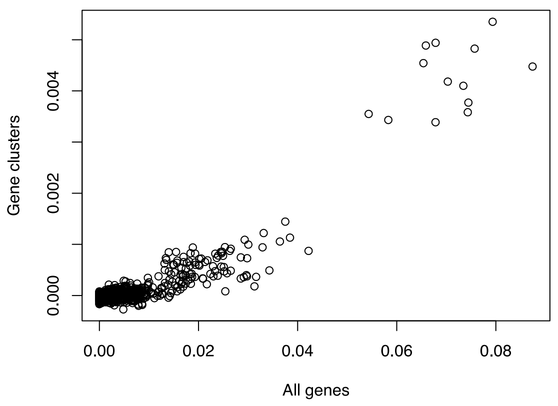


Ranked Spearman MDA, 3500 clusters

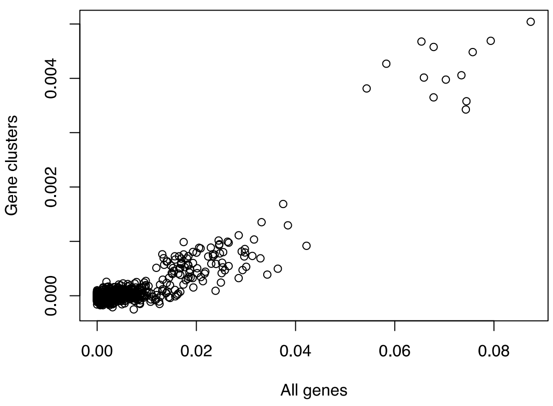


Scaled Euclidean MDA, 4000 clusters

**Random forest importance compared to association testing**

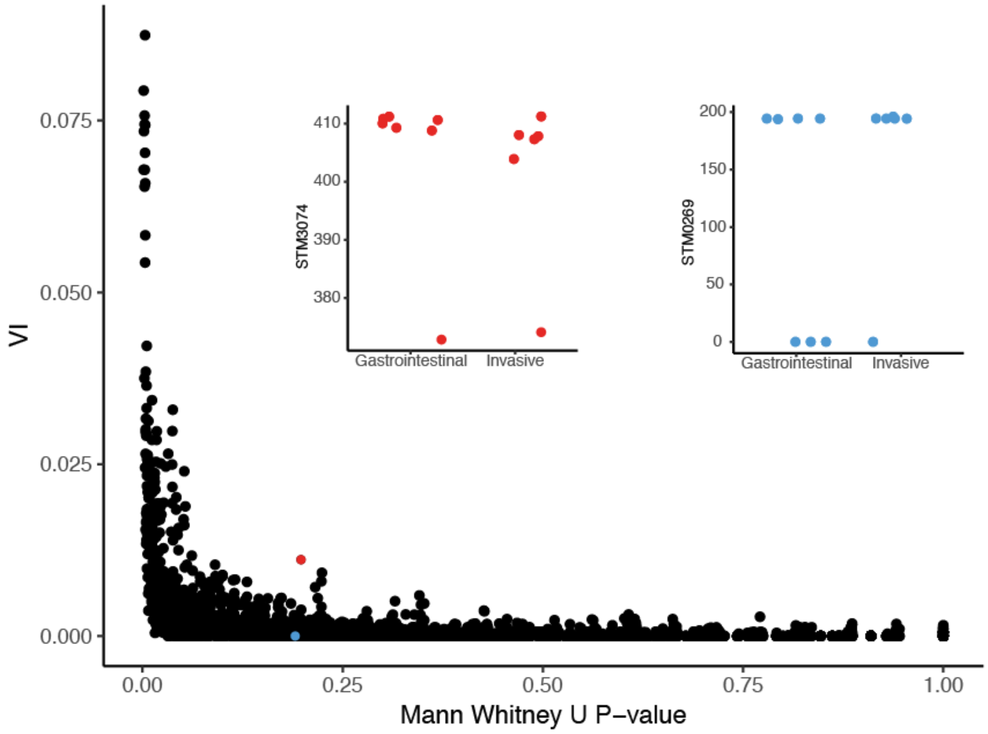


We expect there to be a strong relationship between simple association testing metrics and random forest variable importance (VI), but wanted to explore this more closely to determine whether there was any additional value in using a more complex approach. Correlation between VI and log(Mann Whitney U P-value) was high (Pearson correlation coefficient = -0.57). Note that Mann Whitney U P-values have not been corrected for multiple testing, as this would lead to all P-values approaching ~1, making comparison to VI irrelevant. There were some discrepancies between the two measure, such as that highlighted by the red and blue genes, where Mann Whitney U P-value was similar for a collection of genes, but the distribution of bitscores differed in how useful it was in separating invasive and non-invasive strains when taken in combination with other genes.

In the example case, the red gene appears to consistently accumulate more deleterious mutations than those in gastrointestinal serovars, whereas in the blue gene, there appear to be two sequence variants of the protein, one slightly more common in gastrointestinal serovars than the other.
