## Supplementary Materials for "Machine learning identifies signatures of host adaptation in the bacterial pathogen *Salmonella enterica*"

### Supplemental Material

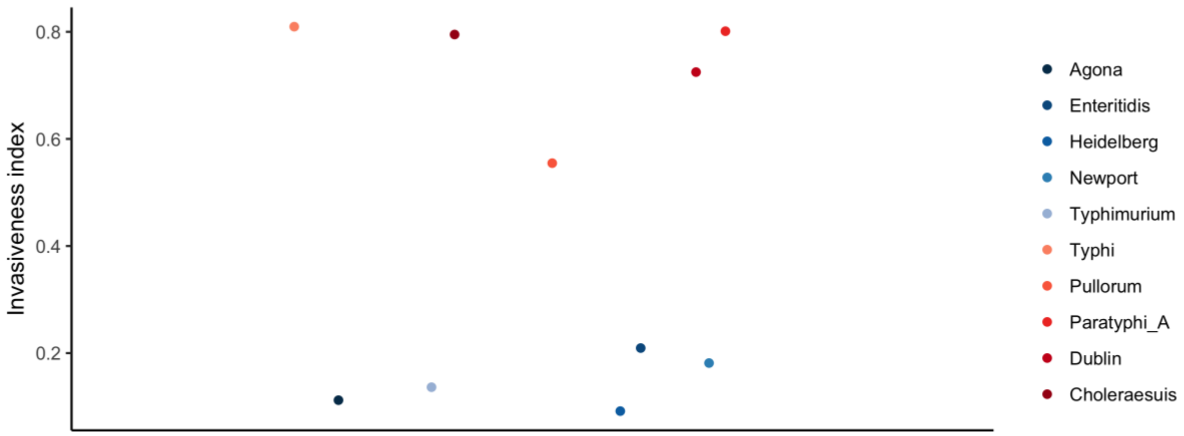


**Figure S1 | Invasiveness index assigned to validation strains of *Salmonella***

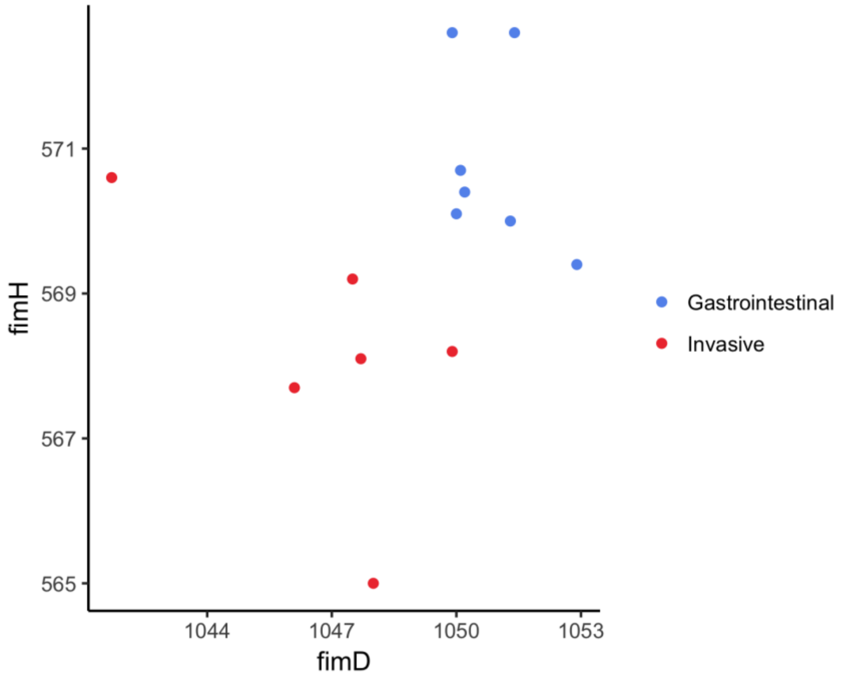


**Figure S2 | Bitscore values from the genes from the genes fimD and fimH combined are better predictors of phenotype than either gene individually**

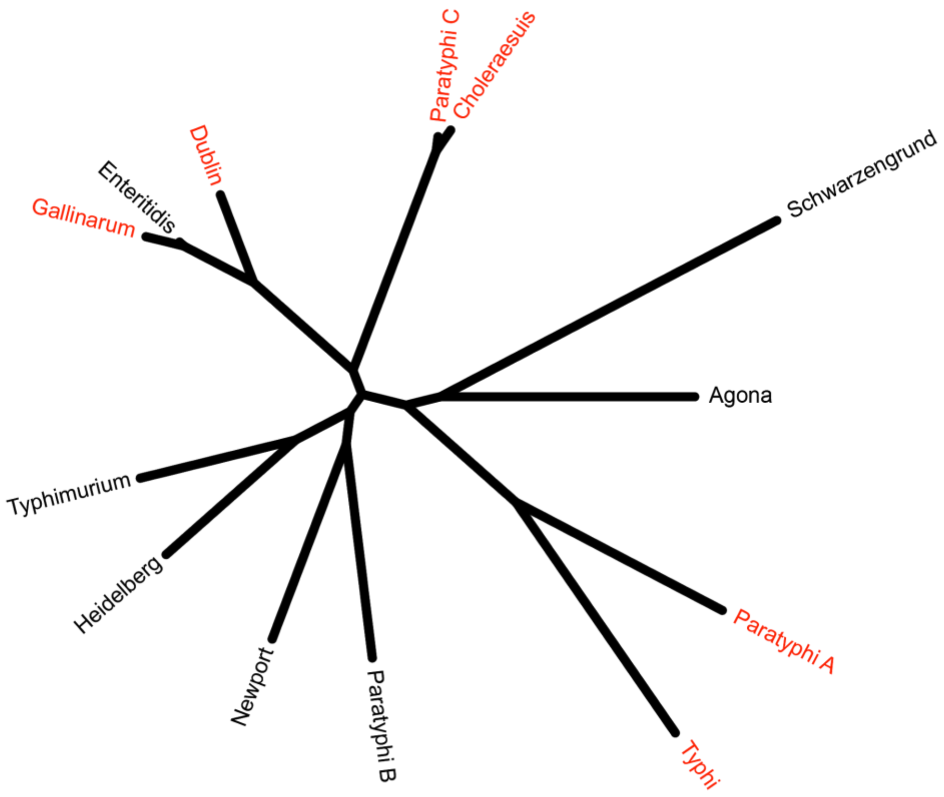


**Figure S3 | Phylogeny of the Salmonella serovars used in this study, constructed in RAxML using a core gene alignment produced by Roary. Invasive serovars are highlighted in red.**

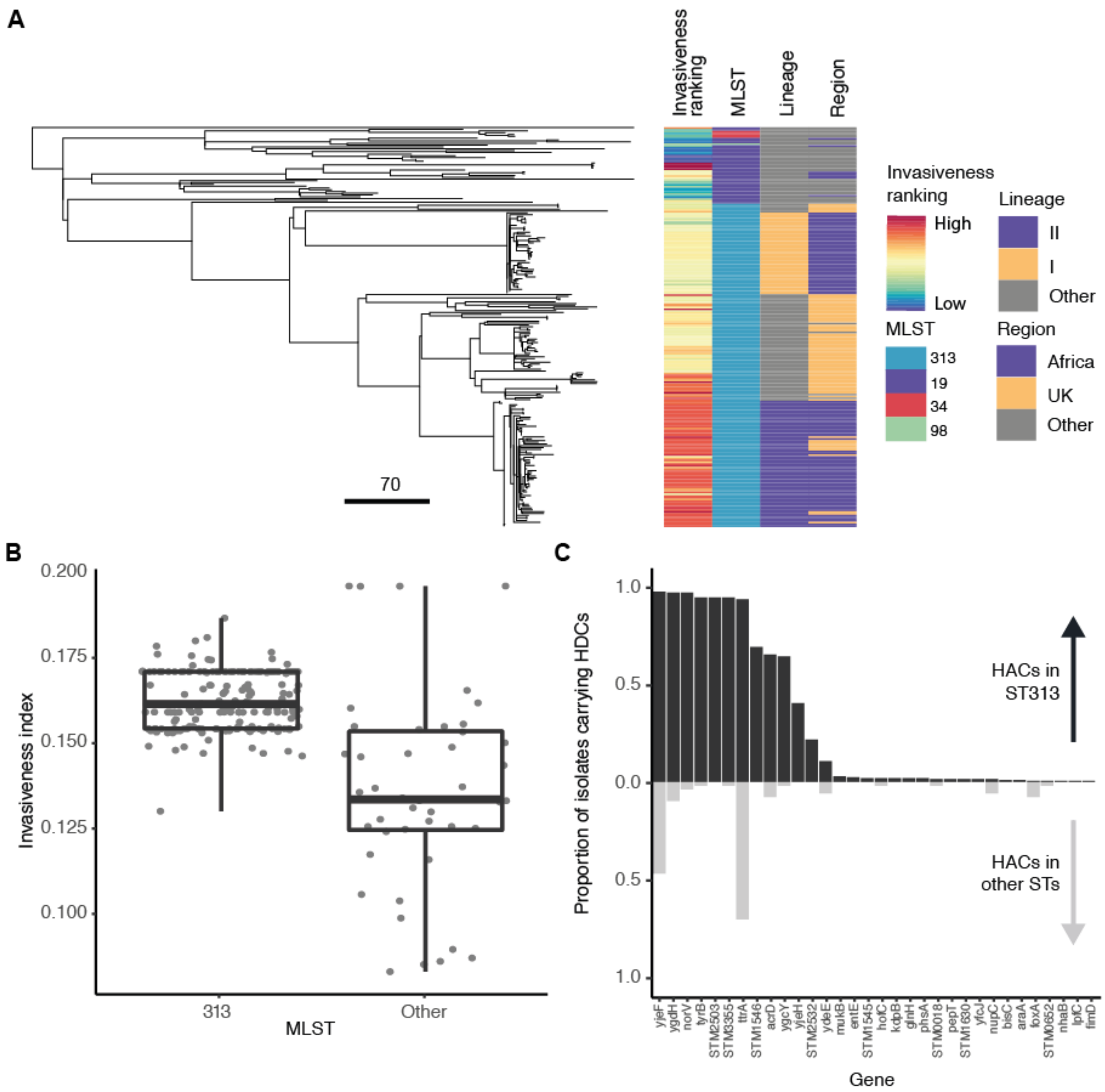


**Figure S4 | Invasiveness scores for *S.* Typhimurium ST313 isolates**

**A: RAxML tree of all ST313 isolates included in the study, annotated with invasiveness ranking and lineage. B: Invasiveness index for all ST313 isolates. C: Proportion of isolates carrying HACs in ST313 compared to other sequence types.**

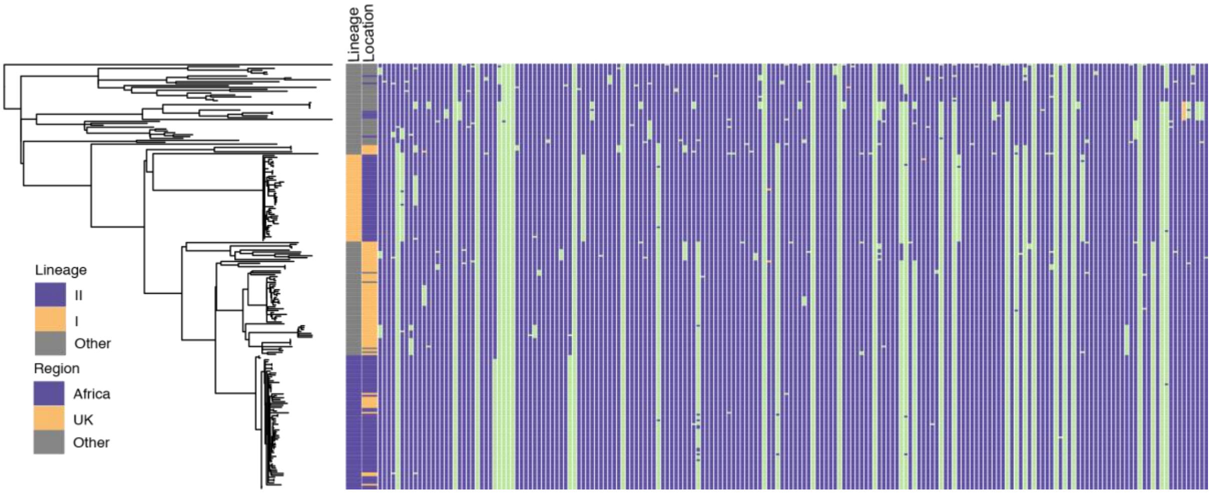


**Figure S5 | Genes in S. Typhimurium ST313 above (intact, purple) or below (attenuated, green) bitscore threshold defined by random forest model for detecting gene degradation associated with invasive isolates. Genes for which homology to the reference sequence was not detected (usually due to extreme truncation) are marked in orange.**

### Supplemental Table S1 | Accession numbers of *Salmonella* enterica serovars used in this study

| Accession | Serovar |
|---|---|
| AE006468 | Typhimurium LT2 |
| CP001113 | Newport SL254 |
| CP001120 | Heidelberg SL476 |
| CP001127 | Schwarzengrund CVM19633 |
| CP001138 | Agona SL483 |
| AM933172 | Enteritidis P125109 |
| CP000886 | Paratyphi B SPB7 |
| AE014613 | Typhi Ty2 |
| FM200053 | Paratyphi A AKU_12601 |
| CP001144 | Dublin CT_02021853 |
| AM933173 | Gallinarum 287/91 |
| AE017220 | Choleraesuis SC-B67 |
| CP000857 | Paratyphi C RKS4594 |
| **Validation strains** | |
| FQ312003 | Typhimurium SL1344 |
| CP007559.1 | Newport CDC2010K-1259 |
| CP016507 | Heidelberg SH12_003 |
| CP006876 | Agona 24249 |
| CP007507 | Enteritidis Durban |
| AL513382 | Typhi CT18 |
| FM200053 | Paratyphi A AKU 12601 |
| CM001151 | Dublin SD3246 |
| CP003786 | Gallinarum bv. Pullorum |
| CM001062 | Choleraesuis A50 |

### Supplemental Table S2 | Top predictor genes

Variable importance: importance assigned to the gene by the random forest model. Wilcox P-value: Wilcox rank sum test for difference in DeltaBS distribution between invasive and gastrointestinal serovars. Non-zero bitscore range: range of bitscores observed across isolates, excluding zeroes assigned to gene deletions and genes which scored too poorly to have a reported match to the HMM for that gene family.

File included separately.

### S3 Table | Counts of high-impact (DBS in the top quartile) independent mutations in each top predictor gene for each strain

File included separately.

### S4 & 5 Tables | Metadata and invasiveness indices for S. Enteritidis and Typhimurium iNTS strains

Files included separately.
